# Beyond ex vivo and in vivo CAR T: antigen-driven CAR T (adCAR-T) expansion method enables rapid, physiological CAR T cells programming

**DOI:** 10.64898/2026.05.15.725377

**Authors:** Andrey Samsonov

## Abstract

Chimeric antigen receptor (CAR) T-cell therapy has demonstrated transformative efficacy in hematologic malignancies, but its broader use remains constrained by complex ex vivo manufacturing, prolonged production timelines, high cost, and dependence on lymphodepleting chemotherapy. Emerging in vivo CAR-T generation strategies aim to address these limitations, but they introduce additional safety concerns associated with systemic delivery of gene-modifying vectors, including off-target transduction and insertional mutagenesis. This paper describes a novel antigen-driven CAR T-cell expansion platform (adCAR-T) based on co-culture of CAR T cells with engineered target cells expressing defined antigen density and lacking the inhibitory checkpoint ligand PD-L1. This system induces immediate activation, rapid proliferation, and sustained cytotoxic differentiation of CAR T cells without reliance on artificial CD3/CD28 bead stimulation or exogenous cytokine-driven expansion. In contrast to conventional methods, the platform eliminates the lag phase of CAR T-cell expansion and enables rapid scaling to clinically relevant doses (10^8^–10^9^ cells) within several days, depending on the initial cell input. Mechanistically, antigen-driven CAR engagement and target-cell lysis trigger cytokine release and amplification of CAR T cells in a physiologically relevant manner. This process promotes coordinated expansion of both directly antigen-engaged and non-engaged CAR T cells. The platform preserves “functional fitness”, minimizes exhaustion, and avoids systemic exposure to gene-delivery vectors. Taken together, this strategy defines a hybrid manufacturing paradigm that bridges the control of ex vivo production with the physiological logic of in vivo activation. Proposed method has a potential to reduce manufacturing complexity, improve safety, and possibly decrease or eliminate the need for lymphodepleting conditioning. This work presents a potential alternative to both standard ex vivo manufacturing and emerging in vivo CAR-T generation approaches, with important implications for improving the accessibility, safety, and cost-effectiveness of CAR T-cell therapies.

## Introduction

Adoptive cell therapy with chimeric antigen receptor (CAR)-engineered T cells has transformed cancer treatment and led to multiple regulatory approvals for B-cell malignancies (Liu et al., 2022). In current autologous CAR-T workflows, T cells are collected from the patient, genetically engineered, expanded ex vivo, subjected to quality control testing, and then returned to the treatment center for infusion. CAR-T therapy is therefore both a biological product and a personalized living therapy manufactured from the patient’s own cells. Despite its remarkable clinical success, broader implementation remains limited by manufacturing complexity, long production timelines, high cost, and restricted scalability. Standard workflows require leukapheresis, T-cell activation using artificial CD3/CD28 stimulation, genetic modification, prolonged ex vivo expansion, and extensive quality testing before reinfusion (Roddie et al., 2019). Because these processes depend on specialized infrastructure and highly controlled manufacturing systems, CAR-T therapy cannot be delivered at every hospital. This creates geographic and institutional disparities in access, forcing some patients to travel long distances or remain near major treatment centers for extended periods. These burdens are not only physical and psychological, but also practical, affecting families, employment, and daily life. In resource-limited settings, reimbursement, regulation, infrastructure, and staffing further constrain implementation.

Traditional CAR-T manufacturing is also expensive and operationally demanding. Specialized facilities generate autologous CAR-T cells by exposing patient-derived T cells to lentiviral vectors that deliver the CAR transgene, followed by expansion to clinically relevant doses. Although the field is now expanding beyond hematologic malignancies into solid tumors, autoimmune diseases, and other indications, the central manufacturing bottleneck remains. Some patients die of progressive disease during the weeks required to manufacture autologous CAR-T products, and others are not eligible because leukapheresis cannot yield sufficient numbers of functional T cells (Qayed et al., 2022).. The cost of treatment, often reaching hundreds of thousands of dollars, places CAR-T therapy beyond the reach of many patients. In addition, standard administration generally requires lymphodepleting chemotherapy before infusion, adding further risk and complexity.

To reduce manufacturing costs and increase throughput, several companies are developing automated CAR-T production platforms (Cellares, 2026). These closed and integrated manufacturing systems aim to increase productivity, reduce batch cost, and lower process failure rates compared with conventional contract manufacturing models. However, even if these platforms succeed commercially, they will still produce CAR-T cells using fundamentally similar ex vivo logic and are therefore unlikely to solve the biological limitations associated with current manufacturing paradigms. A key limitation of conventional expansion is the reliance on non-physiological activation signals, which may promote T-cell exhaustion and diminish in vivo persistence. Prolonged culture further exacerbates functional decline, as reflected by increased exhaustion markers and reduced cytotoxic capacity over time. Clinical experience has shown that the quality and phenotype of ex vivo-expanded CAR-T cells can strongly influence therapeutic efficacy.

The first CAR-T cell therapy approved by the U.S. Food and Drug Administration was tisagenlecleucel (Kymriah, Novartis), approved in 2017 for children and young adults with relapsed or refractory B-cell precursor acute lymphoblastic leukemia. It was also the first gene therapy approved in the United States. Its approval was based on a high remission rate in a pivotal clinical trial. However, product characterization revealed substantial lot-to-lot variability in functional potency, including marked differences in interferon-γ (IFN-γ) production in response to antigen-bearing target cells (Fig. 1).

**Figure 1.**
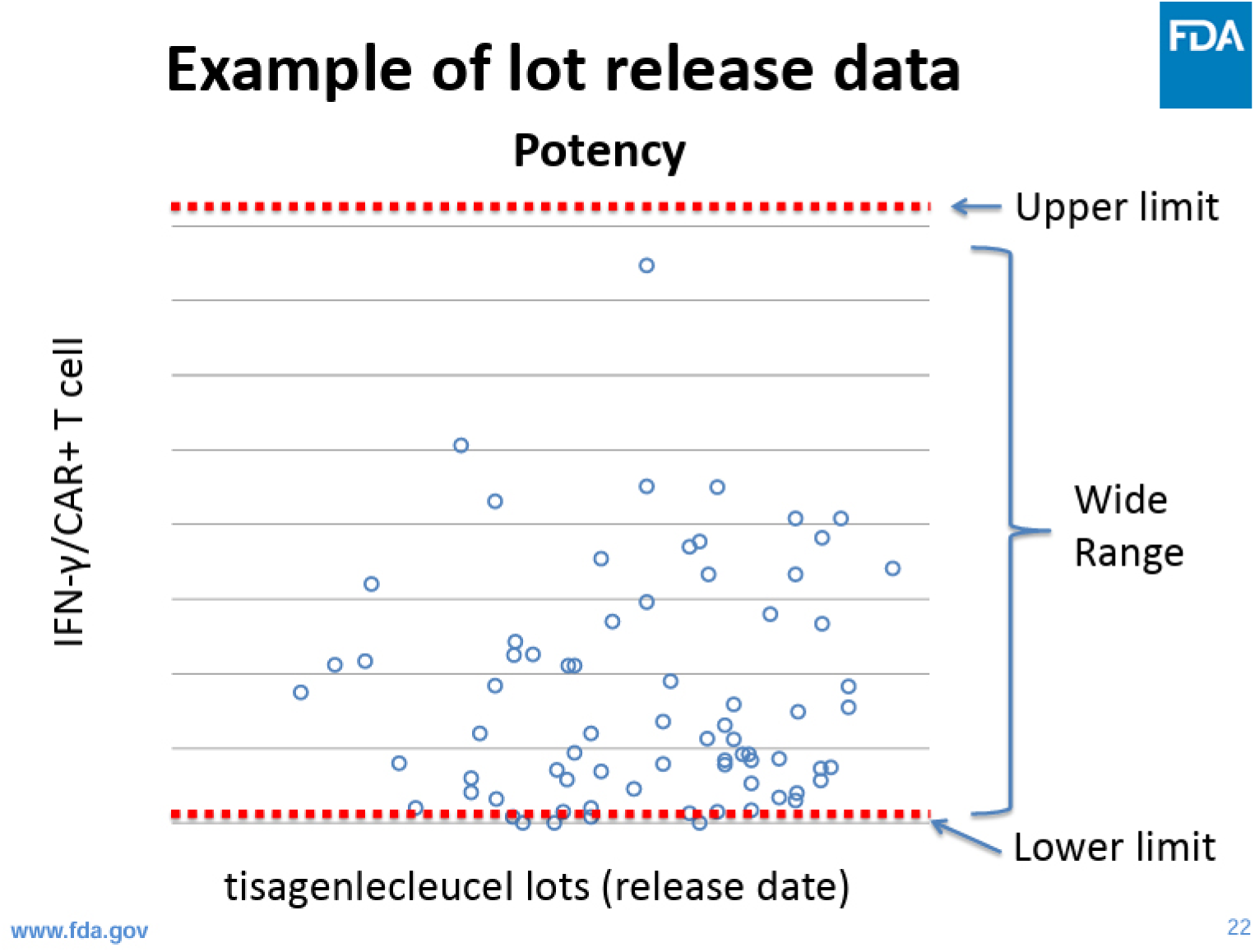
INF-γ production per transduced cell. INF-γ produced during co-culture with CD19-expressing cells (FDA generated) (Meeting, 2017).

Because IFN-γ release is widely used as a surrogate for CAR-T activation and potency, this variability made it difficult to correlate in vitro measurements with clinical safety or efficacy. Likewise, the percentage of CAR-positive T cells varied across manufactured lots (Fig. 2). Since the active component of the product is the CAR-expressing T-cell fraction, control of the transduction step is a central manufacturing concern. In practice, this attribute is typically regulated by adjusting the amount of vector used per cell during transduction, commonly expressed as multiplicity of infection. Yet these examples highlight a broader issue: the ultimate goal of CAR introduction into autologous T cells is not merely successful transduction in vitro, but effective tumor recognition and elimination in the patient.

**Figure 2.**
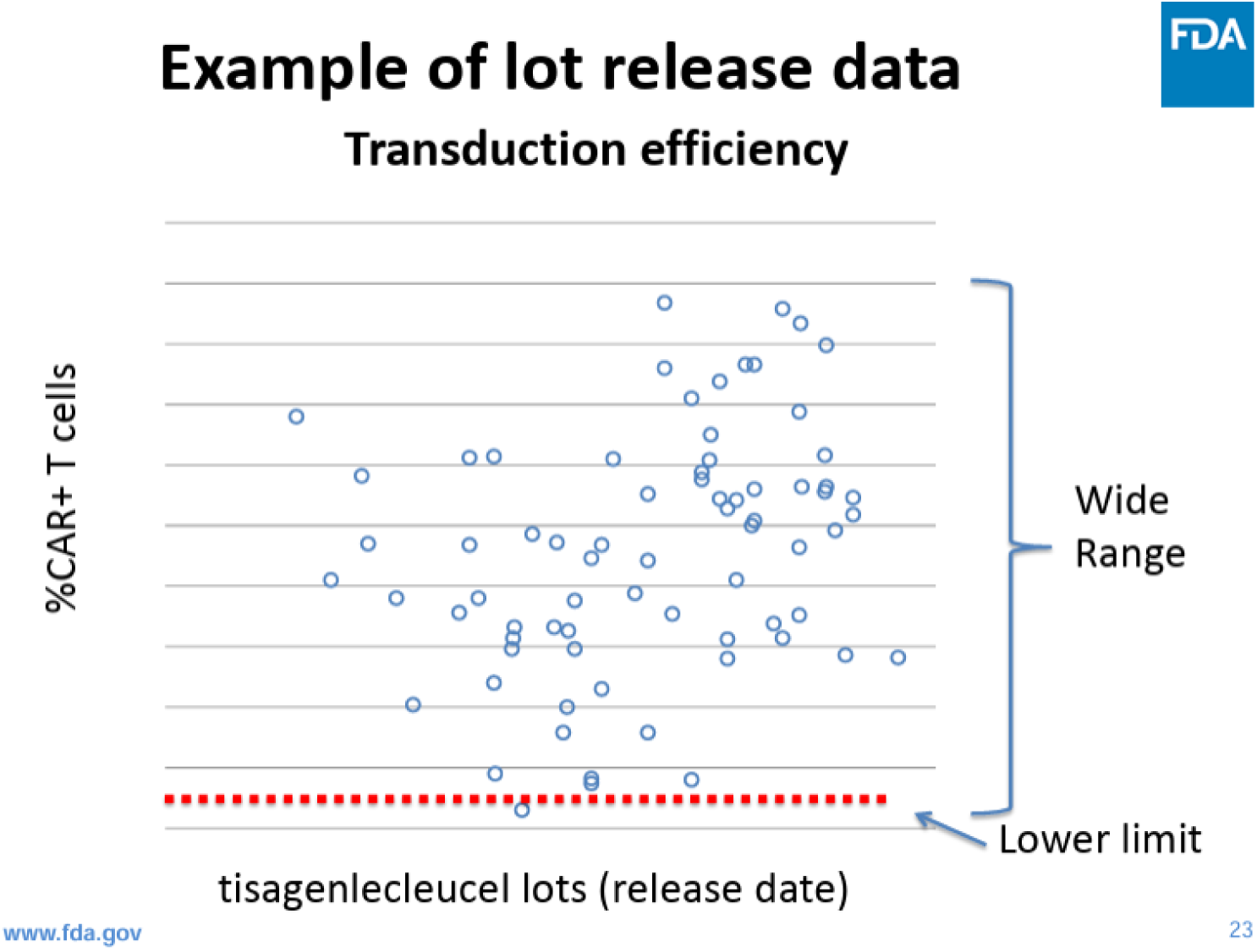
Results of percent CAR-positive T cells in tisagenlecleucel. (FDA generated) (Meeting, 2017).

Because the active ingredient of the product is CAR-positive T cells, transduction efficiency is used to calculate the final cell dose (number of CAR-positive cells). Therefore, in order to produce tisagenlecleucel with consistent product quality attributes, it is essential to control the transduction step. This product attribute is typically controlled through the amount of vector used per cell during transduction i.e. the multiplicity of infection (MOI).

Main FDA concern was that in the clinical trials, IFN-γ production and percentage of CAR T positive cells varied greatly from CAR-T lot-to-lot (Fig. 1, 2), making it difficult to correlate IFN-γ production *in vitro* to CAR-T expressing T cells efficacy in patients. Indeed, main goal of CAR T introduction in autologous patient T cells is effective hunting for tumor cells inside of human body.

Magnetic bead-based activation remains the dominant approach for ex vivo T-cell expansion, although optimal activation and expansion conditions have not been fully defined. In standard methods, T cells are stimulated with magnetic polymer beads coated with monoclonal antibodies against CD3 and CD28. Less commonly, anti-CD3 and anti-CD28 antibodies are immobilized directly onto culture vessels. As an alternative, colloidal nanomatrix systems conjugated to anti-CD3 and anti-CD28 antibodies have also been developed (TransAct™), with the advantage that excess activation reagent can be removed more easily by centrifugation or medium exchange (Biotec, 2026).. However, growth curves of T cells activated with CD3/CD28 reagents and interleukin-2 often show a long lag phase before proliferation begins, sometimes lasting 7–12 days (Fig. 3). Only after this delay do cells enter a period of rapid expansion. Thus, the combined activation and expansion process may require around 20 days, contributing substantially to the time and cost of production.

**Figure 3.**
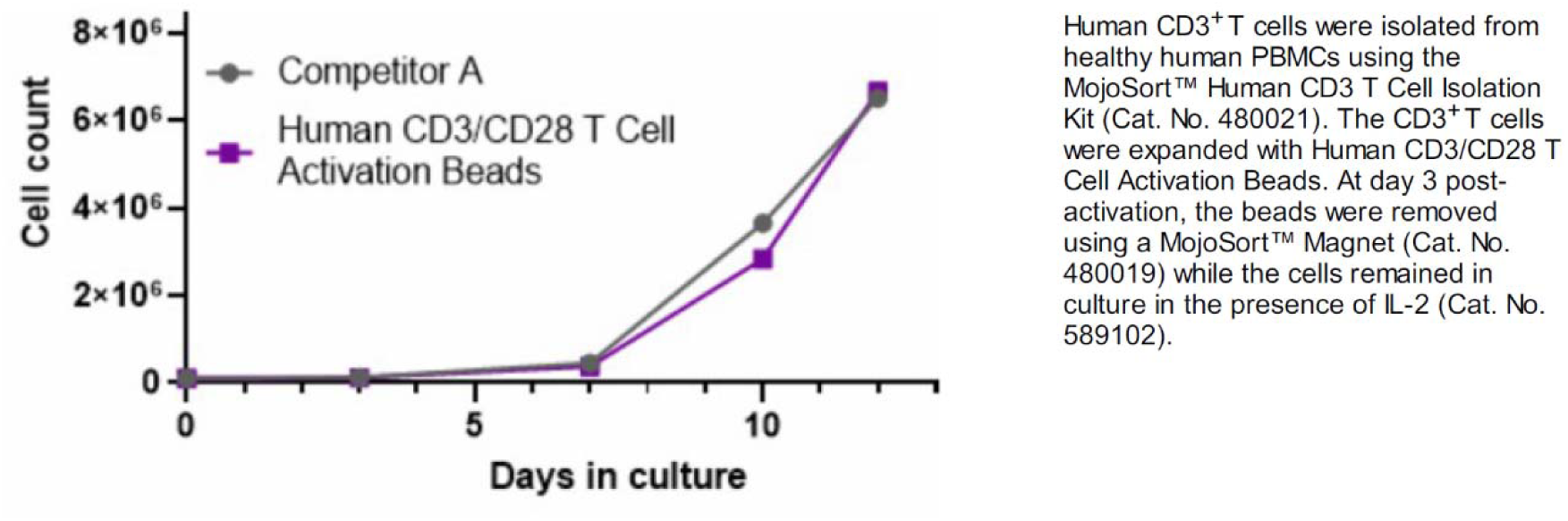
Human CD3 T cells were isolated from healthy human PBMCs using the MojoSort™ Human CD3 T Cell Isolation Kit (Cat. No. 480021). The CD3 T cells were expanded with Human CD3/CD28 T Cell Activation Beads. At day 3 post activation, the beads were removed using a MojoSort™ Magnet (Cat. No. 480019) while the cells remained in culture in the presence of IL-2 (Cat. No. 589102) (BioLegend, 2026).

Cell-based artificial antigen-presenting cells (aAPCs) offer a more physiological alternative. In one study, CAR-T cells were expanded either with anti-CD3/CD28-coated beads or with engineered K562-based aAPCs expressing CD19, CD64, CD86, CD137L, and membrane-bound IL-15 than that with beads (Yang et al., 2021). The aAPC platform yielded greater expansion of CD19-specific CAR-T cells than the bead-based method, along with higher representation of memory phenotypes and lower exhaustion. In addition, aAPC-expanded CAR-T cells retained strong antitumor activity in vitro and in vivo, and secreted substantial amounts of cytokines after tumor encounter, suggesting preserved function. These findings indicate that cell-based stimulation can outperform bead-based activation in relevant settings. Notably, according to literature K562 cells generally express low basal levels of PD-L1, and IFN-γ stimulation increases this expression only modestly (see Fig. 1A from (Berthon et al., 2010)). However, despite these promising results, the authors did not propose a manufacturing strategy centered on serial co-culture of CAR-T cells with engineered target cells as a dedicated expansion platform.

Another important effort to improve access has been the development of allogeneic CAR-T therapies. These “off-the-shelf-like” approaches use donor-derived cells or similar sources rather than manufacturing a separate product for each patient. Compared with autologous CAR-T therapy, allogeneic models may offer faster availability and greater manufacturing standardization. They also bring their own challenges, including rejection, graft-versus-host disease, and limited persistence. Nonetheless, they reflect the same broader goal as in vivo CAR-T strategies: reducing the burden of individualized ex vivo manufacturing (Lonez and Breman, 2024).

Rapid-manufacturing approaches have also emerged as alternatives to prolonged ex vivo expansion. So-called “next-day” CAR-T methods aim to generate functional, minimally activated CAR-T products within 24 hours (Ghassemi et al., 2022). The “next-day” (24-hour) protocols generate functional CAR T cells from obtained patient T cells within one day (Yang et al., 2022) or during 3 days for FasTCAR (Gracell, 2026). In mouse xenograft models, rapidly generated non-activated CAR-T cells have shown greater antileukemic activity per cell than conventionally expanded CAR-T cells (Ghassemi et al., 2022), suggesting that shorter ex vivo manipulation may preserve functional potency. Several platforms (like FasTCAR, InstanCAR-T) claim that these rapidly manufactured products can be infused without extensive ex vivo expansion because the cells remain highly proliferative and cytotoxic after infusion. Early clinical results have been encouraging with some patients achieving remission in 14 days, but they remain in the research and trial phase (Zhang et al., 2022) and are not yet FDA approved.

In parallel, in vivo CAR-T generation strategies have gained substantial attention. These approaches seek to bypass ex vivo manufacturing process (lasts on average 14–21 days) that typically delivers between 10 million and 250 million CAR T-cells (Pierini et al., 2026). In vivo CAR T-cell therapy is delivering gene-modifying constructs, such as lentiviral (LV) vectors or mRNA-loaded lipid nanoparticles (LNPs), directly to immune cells in the patient. Conceptually, in vivo CAR-T could transform the field by eliminating or reducing apheresis, individualized manufacturing, and complex logistics (Pierini et al., 2026). If successful, such strategies could shorten treatment timelines, reduce cost, and make CAR-T therapy more accessible beyond major academic centers (Nyberg et al., 2026). This promise has generated considerable enthusiasm and even some euphoria. In vivo CAR T enthusiasts state: “If we can translate this to humans, we could dramatically reduce costs, eliminate waiting times, and potentially allow community hospitals — not just major cancer centers. That would truly democratize access to CAR-T cell therapy.” The recent clinical reports suggest that in vivo CAR-T generation is becoming technically feasible. However, clinical experience remains limited, and safety concerns remain substantial. Interius BioTherapeutics announced first-patient dosing with its in vivo CAR gene therapy and later reported expansion of clinical permissions in Europe (Interius, 2024). Umoja Biopharma received FDA Fast Track designation for its in vivo CAR-T candidate for CD19-positive B-cell malignancies and had previously announced IND clearance in 2024 (Umoja, 2025). In addition, a Nature Medicine paper published on March 25, 2026 reported a phase 1 study of anti-BCMA in vivo CAR-T in five patients with relapsed or refractory multiple myeloma, with objective responses in 4 of 5 patients and stringent complete remission in 3 of them (An et al., 2026). ESO-T01 is a nanobody-directed, immune-shielded lentiviral vector encoding a humanized anti-B cell maturation antigen (BCMA) CAR, in adults with relapsed or refractory multiple myeloma (An et al., 2026). ESO-T01 was administered as a single intravenous infusion of 0.2 × 10^9^ transduction units (TU) without leukapheresis, ex vivo manufacturing or lymphodepleting chemotherapy. The cohort is very small and follow-up remains limited, so broad conclusions would be premature. Still, it is increasingly clear that the idea of “generating CAR-T inside the body” is beginning to look clinically real. These findings provide preliminary evidence on the feasibility and safety of in vivo CAR-T generation using an immune-shielded vector. While conceptually attractive, the in vivo strategies introduce new risks, including uncontrolled biodistribution, off-target transduction, and insertional mutagenesis associated with viral vectors. Many clinical observations highlight that such risks remain non-negligible despite improvements in vector design.

Intravenous administration of lentiviral vectors introduces the possibility of uncontrolled biodistribution, off-target transduction and insertional mutagenesis. LV vectors integrate sequences permanently into host cell chromatin using viral integrase. LV integration into cell genome can induce alternative splicing, producing aberrant, chimeric transcripts by utilizing cryptic splice sites within the vector’s long terminal repeat (LTR) or gag gene (Moiani et al., 2012). These aberrant transcripts are often recognized and eliminated by cellular mechanisms like nonsense-mediated mRNA decay (NMD) and other cell quality control mechanisms (Suwanmanee et al., 2017). The genotoxicity risks of LVs are mainly related to aberrant transcriptional activation or inactivation of cellular genes and the induction of new splice variants with potentially oncogenic effects (Cavazza et al., 2013).

Confirming these facts, clinical experience with lentiviral gene therapies in other settings has shown that secondary hematologic malignancies can emerge months to years after treatment, driven by vector integration into proto-oncogenes or disruption of tumor suppressor pathways (Mast, 2024). The study published in *The New England Journal of Medicine (NEJM)* show that seven out of 67 patients under 18 years of age who took part in a Phase II and Phase III trial for Skysona developed haematologic cancers (Duncan et al., 2024). The patients received the one-time autologous hematopoietic stem cell-based gene therapy as a treatment for early cerebral adrenoleukodystrophy (CALD) – a rare and fatal neurodegenerative disease. Researchers at Boston Children’s Hospital analyzed blood and bone marrow samples from patients in the clinical studies ALD-102 (NCT01896102) and ALD-104 (NCT03852498), along with an ongoing follow-up study called LTF-304. The researchers found one patient with haematological cancer from ALD-102, whilst six came from the ALD-104 study. Myelodysplastic syndrome (MDS) accounted for six of the cases, whilst one patient had acute myeloid leukaemia (AML). Occurrences of blood cancer ranged from 14 months after therapy to around 7.5 years following Skysona treatment. The observed cases of haematologic malignancy appear to be driven by integration of Skysona’s lentiviral vector into a proto-oncogene, as per an FDA analysis document during the therapy’s approval, and the NEJM publication’s authors. These observations indicate that genotoxic risk remains real, even with improved vector design. Accordingly, in vivo CAR-T strategies that rely on integrating viral vectors may inherit these liabilities.

Non-viral in vivo approaches based on lipid nanoparticles loaded with mRNA encoding CAR constructs may appear safer because the delivered mRNA does not integrate into the genome (Billingsley et al., 2020).. In these systems, transient CAR expressions can in principle enhance safety, support repeat dosing, and permit greater pharmacologic control. Yet lipid nanoparticles also carry their own liabilities. Clinical experience with mRNA-LNP vaccines and therapeutics has shown that these formulations can trigger acute dose-dependent inflammatory toxicities, including local and systemic reactions, cytokine release, and, in some settings, myocarditis (Kadali et al., 2021; Wu and Wang, 2025). Although improvements in lipid chemistry, targeting ligands, and shielding strategies may reduce off-target biodistribution and inflammatory effects, the safety and efficiency of T-cell-specific in vivo delivery remain unresolved.

A further challenge for in vivo CAR-T approaches is predictability. In standard ex vivo CAR-T therapy, products are typically administered at doses ranging from tens of millions to up to one billion CAR-T cells. By contrast, it is difficult to predict how many functional CAR-T cells will be generated after in vivo delivery of lentiviral vectors or mRNA nanoparticles. Preclinical studies in mice have generated CAR-T cell numbers (1 to 5 million CAR-T cells) sufficient for efficacy in murine models, but these scales are not directly comparable to human treatment requirements (Zhao et al., 2020). Moreover, in vivo approaches must address the possibility that non-T-cell populations, including malignant B cells, may be transduced (Ruella et al., 2018). Such events are not merely theoretical: CAR transgene transfer into a leukemic B cell during ex vivo manufacturing has previously produced a CAR-expressing leukemia clone that became invisible to CAR-T recognition through cis masking of CD19, ultimately contributing to fatal disease progression (Ruella et al., 2018). Similar events could, in principle, occur with in vivo delivery, where control over target-cell specificity is even more difficult. The core questions therefore remain unresolved: can off-target transduction be sufficiently prevented, can expression be controlled tightly enough, and can toxicities such as cytokine release syndrome and neurotoxicity be safely managed in an in vivo setting? These are not minor engineering details, but central regulatory and clinical challenges.

Here we describe an antigen-driven CAR-T expansion strategy designed to accelerate manufacturing while preserving functional potency and resistance to exhaustion and senescence. We refer to this approach as the adCAR-T or “battlefield” method, because the cells expand through repeated cycles of antigen encounter and killing. T cells, including CAR-T cells, are known “serial killers”: a single cytotoxic cell can eliminate many target cells (up to 20) before eventually becoming exhausted or dying (Davenport et al., 2015; Liadi et al., 2015). In standard ex vivo manufacturing, however, CAR-T cells never encounter their intended targets before infusion. By contrast, the adCAR-T method uses engineered target cells to drive repeated, physiologically relevant CAR engagement during expansion. This creates a self-sustaining amplification process in which expanding CAR-T cells release endogenous cytokines that support their own survival, persistence, and continued proliferation.

The continued presence of engineered target cells may also provide selective cues that better prepare CAR-T cells for the in vivo tumor environment. Tumor cells are known to secrete chemokines such as CXCL9, CXCL10, and CXCL11, which attract and stimulate antitumor CD8+ T cells, particularly under inflammatory conditions (Abdul-Rahman et al., 2024; Ozga et al., 2021). It is therefore plausible that the adCAR-T method favors expansion of CAR-T cells better adapted to recognize such chemotactic signals and to localize to sites of high tumor burden. In this sense, antigen-driven expansion may not only accelerate manufacturing but also better align the resulting product with the biological demands of tumor control in vivo.

## Results

### Concept and design of the antigen-driven CAR T-cell expansion platform (adCAR-T)

The proposed antigen-driven CAR T-cell expansion platform replaces artificial activation signals, such as beads coated with anti-CD3 and anti-CD28 antibodies, with direct antigen-mediated stimulation using engineered target cells. In the intended workflow, T cells obtained after leukapheresis, and isolation are immediately genetically modified to express a CAR construct. The resulting CAR T cells are then co-cultured with engineered target cells that (i) express the relevant target antigen at a defined surface density, (ii) lack inhibitory checkpoint ligands such as PD-L1, and (iii) must be mitotically inactivated, for example by γ-irradiation. This design enables controlled antigen exposure while preventing uncontrolled target-cell outgrowth. Upon co-culture, CAR T cells rapidly recognize antigen, form immunological synapses (IS), and initiate cytotoxic activity. Target-cell lysis is accompanied by the release of cytokines, including IFN-γ, TNF-α, IL-10, and IL-8, which support both autocrine and paracrine amplification. Importantly, proliferation is not restricted to directly antigen-engaged CAR T cells. Instead, cytokine-mediated signaling promotes coordinated expansion across the broader CAR T-cell population, enabling rapid scaling without the need for artificial CD3/CD28 stimulation. Because IL-10 has been implicated in promoting CAR T-cell expansion, stem-like memory features, and resistance to immunosuppression, its induction may be particularly relevant in this system (Huang and Wang, 2024).

### Immediate activation without a lag phase

A major feature of the adCAR-T platform is the absence of the prolonged lag phase typically observed in conventional CD3/CD28-based expansion systems. Standard ex vivo activation methods often require 7–12 days before robust proliferation begins (Fig. 3). In contrast, antigen-driven stimulation produces immediate functional engagement and early proliferation, markedly shortening the time required to generate large cell numbers.

### Antigen surface density is a critical determinant of expansion

We identify target-antigen surface density as a key parameter governing CAR T-cell activation, proliferation, and functional stability. Prior work has shown that CAR T cells can extract target antigen from tumor cells through trogocytosis, a process that lowers antigen density on tumor cells while transferring antigen to the CAR T-cell surface. This can promote fratricide, functional exhaustion, and ultimately antigen-low relapse (Hamieh et al., 2019).. These effects have been observed across multiple CAR targets, including CD19, CD22, BCMA, and mesothelin, indicating that trogocytosis is a general feature of CAR T-cell biology.

These observations suggest that antigen density must be balanced carefully in the proposed “battlefield” expansion format. If target cells express too little antigen, CAR T cells may fail to form productive immunological synapses and lyse targets inefficiently. Conversely, if antigen density is too high, CAR T cells may form overly stable conjugates that increase trogocytosis, fratricide, and exhaustion. We therefore propose that an intermediate antigen density is optimal, as it minimizes trogocytosis and fratricide while preserving strong target-cell lysis and cytokine production. In this setting, robust lysis is accompanied by the release of IFN-γ, TNF-α, IL-10, and IL-8, which then support expansion of both directly engaged and non-engaged CAR T cells. This coordinated amplification may help replenish CAR T cells lost through repeated killing events.

### Checkpoint-independent target-cell design

Checkpoint signaling is an additional variable that can shape CAR T-cell expansion and persistence. IFN-γ, while beneficial for CAR T-cell activation and amplification, can also act as a double-edged sword by inducing PD-L1 expression on target cells, thereby promoting PD-1-mediated dysfunction and apoptosis in therapeutic T cells (Patsoukis et al., 2020). Although PD-1/PD-L1 blockade can restore function in some settings, complete elimination of PD-1 signaling is not necessarily advantageous, as prior work suggests that excessive activation in the absence of inhibitory signaling can impair long-term survival (Kalinin et al., 2021). These findings indicate that productive CAR T-cell responses require a balance between activation and negative regulation.

To minimize suppressive signaling during ex vivo expansion, we selected the RL cell line (ATCC CRL-2261), a B-lymphoblast line derived from non-Hodgkin lymphoma, because it exhibits low basal PD-L1 expression and does not appreciably upregulate PD-L1 after IFN-γ stimulation. Flow cytometric and microscopy-based analyses confirmed the absence of PD-L1 induction in both wild-type RL cells and the corresponding CD19-knockout derivative after IFN-γ exposure (Fig. 4). We therefore used RL-derived target cells as a checkpoint-silent platform for antigen-driven CAR T-cell expansion.

**Figure 4.**
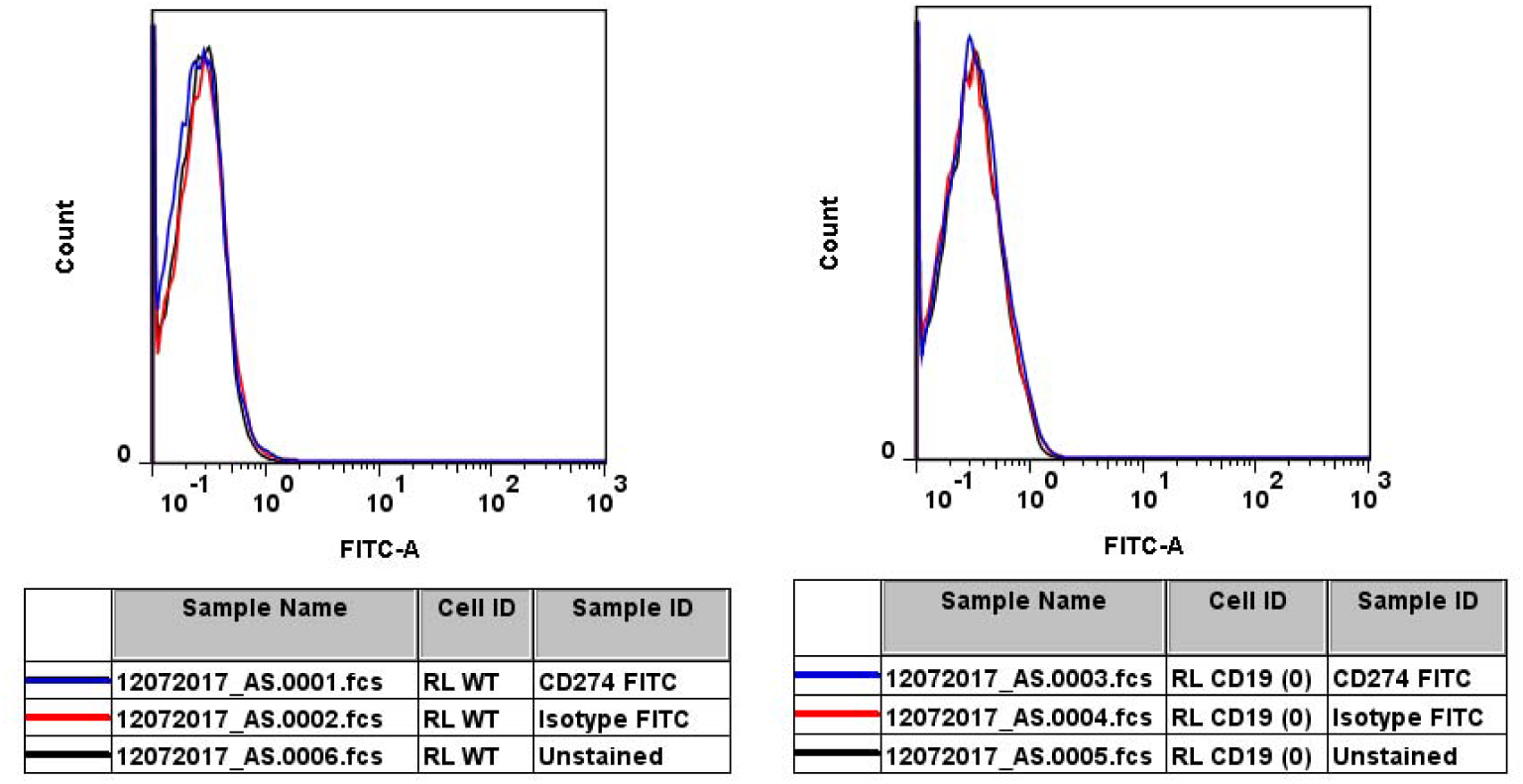
Left: WT RL cells were stimulated with INF-γ and labelled with. Anti-PD-L1 (CD274) antibodies after 20h. Right: RL CD19 KO cells were stimulated with INF-γ and labelled with anti-PD-L1 (CD274) antibodies after 24h. Both WT RL and RL CD19 KO do not express PD-L1 on their surface before and after INF-gamma stimulations.

### Generation of RL target-cell lines with graded CD19 density

To systematically investigate the effect of antigen density, we generated a panel of RL-derived target-cell lines expressing graded levels of CD19. First, a homozygous CD19 knockout clone was generated from wild-type RL cells. During clone selection, a population with intermediate CD19 surface density was identified and found to be heterozygous for CD19 disruption; this pool was retained as an RL CD19 heterozygous knockout population. To create additional defined-expression states, CD19 was reintroduced into the knockout background under the control of either the LMNB1 promoter or the ACTB promoter. These manipulations yielded a panel of five RL-derived cell lines spanning a range of CD19 surface densities, from complete absence of CD19 to very high expression. Flow cytometry and fluorescence microscopy confirmed stable and distinct expression profiles across the panel (Fig. 5). Introduction of CD19 through endogenous-style knock-in strategies offered two conceptual advantages over lentiviral transduction: greater stability of expression and reduced risk of aberrant transcriptional effects associated with random vector integration. Because integrating lentiviral vectors can generate aberrant splice products and other unintended genomic consequences, controlled genomic insertion under defined promoter contexts may be preferable for generating stable target-cell lines used in the expansion platform.

**Figure 5.**
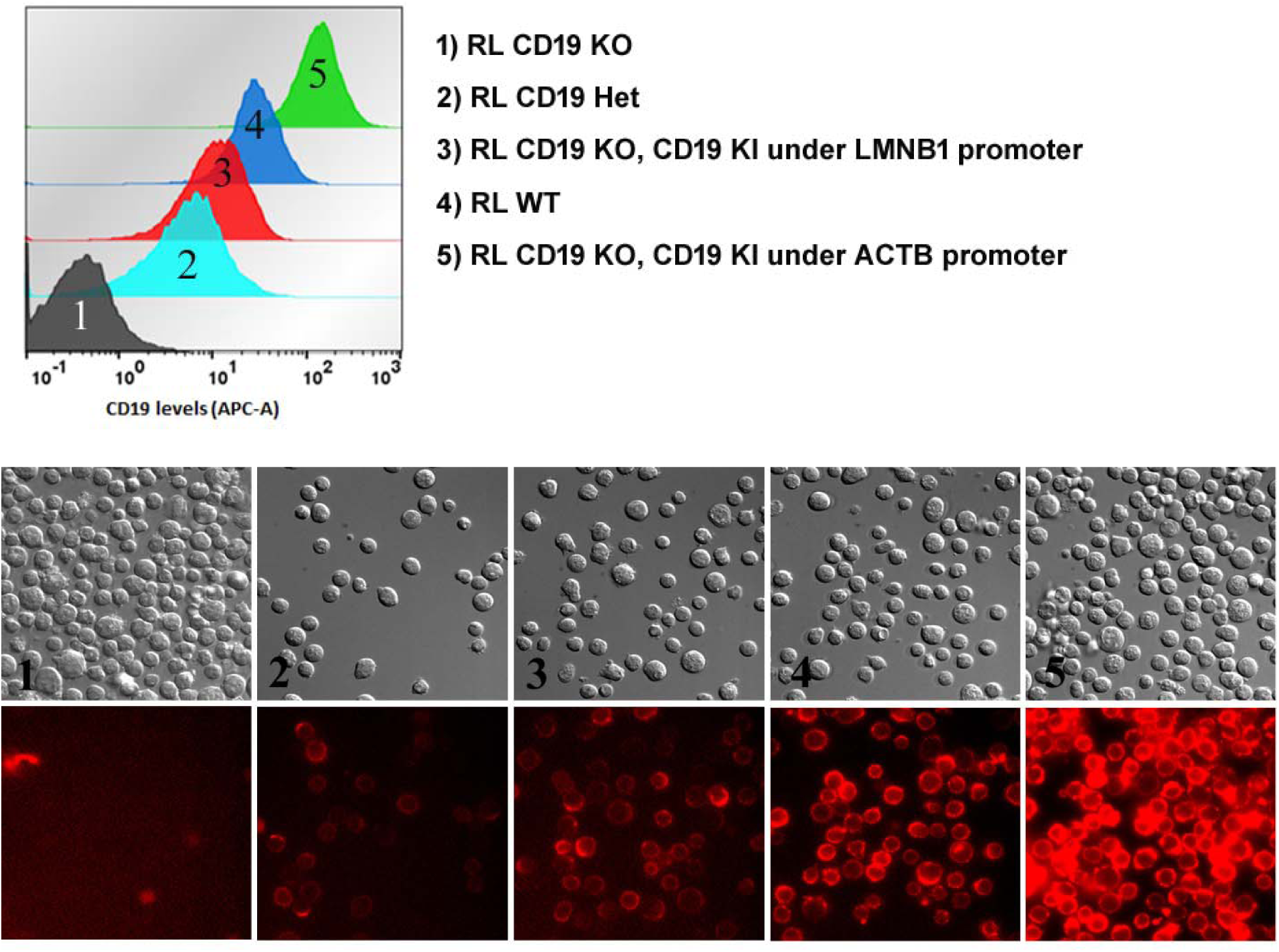
Top: FACS data characterizing the RL CD19 “CAR-T target” cell lines. CD19 gene was KO in RL WT cells and re-introduced under different strength promoters (LMNB1 and ACTB). RL heterozygous CD19 knockout pool was also selected to have low density of CD19. Bottom: immunofluorescence microscopy of CAR T lines (magnification 40x).

**Figure 6.**
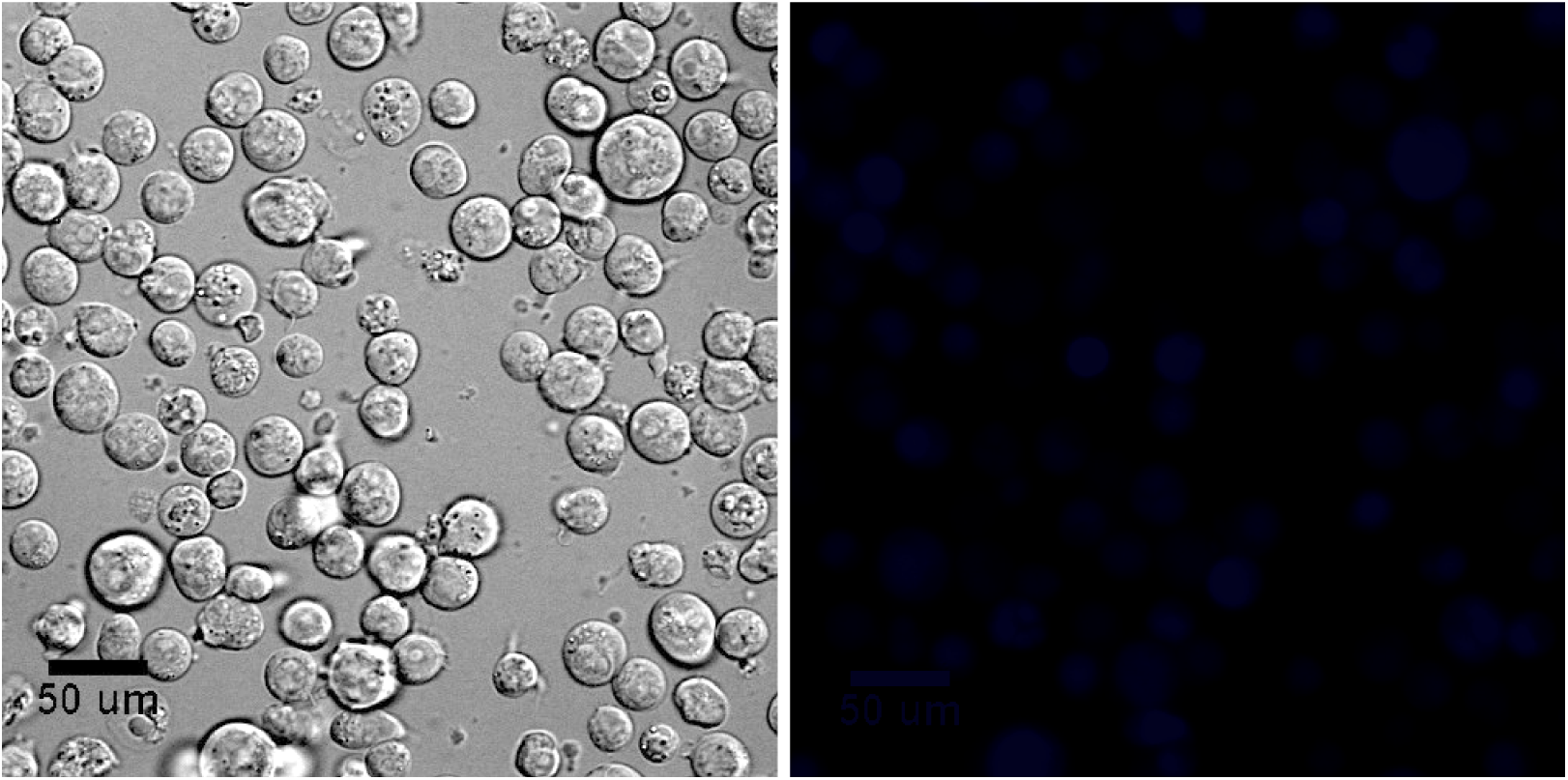
RL cells with BFP viability marker.

For more robust observation of target cells lysis and control of residue target cells during CAR T expanding, it is possible to add viability market to chosen target cell line via genome editing method. For example, Blue Fluorescence Protein (BFP) (Fiig. 6). After CAR T formed immunological synapse and inject Granzyme through Perforin inside of target cell cytoplasm this cell will die via induce programmed cell death (apoptosis).

### CAR T cells fail to expand in the absence of target antigen

When CAR T cells were mixed with RL CD19-knockout cells, no meaningful expansion was observed. In these cultures, the target cells continued to proliferate and formed large aggregates, indicating that CAR T cells could not initiate productive activation in the absence of cognate antigen (Fig. 7). By contrast, when CAR T cells were co-cultured with RL cells expressing CD19 at any detectable level, they formed immunological synapses, released cytokines, and lysed the target cells. These observations confirm that antigen recognition is required to initiate the adCAR-T amplification program.

**Figure 7.**
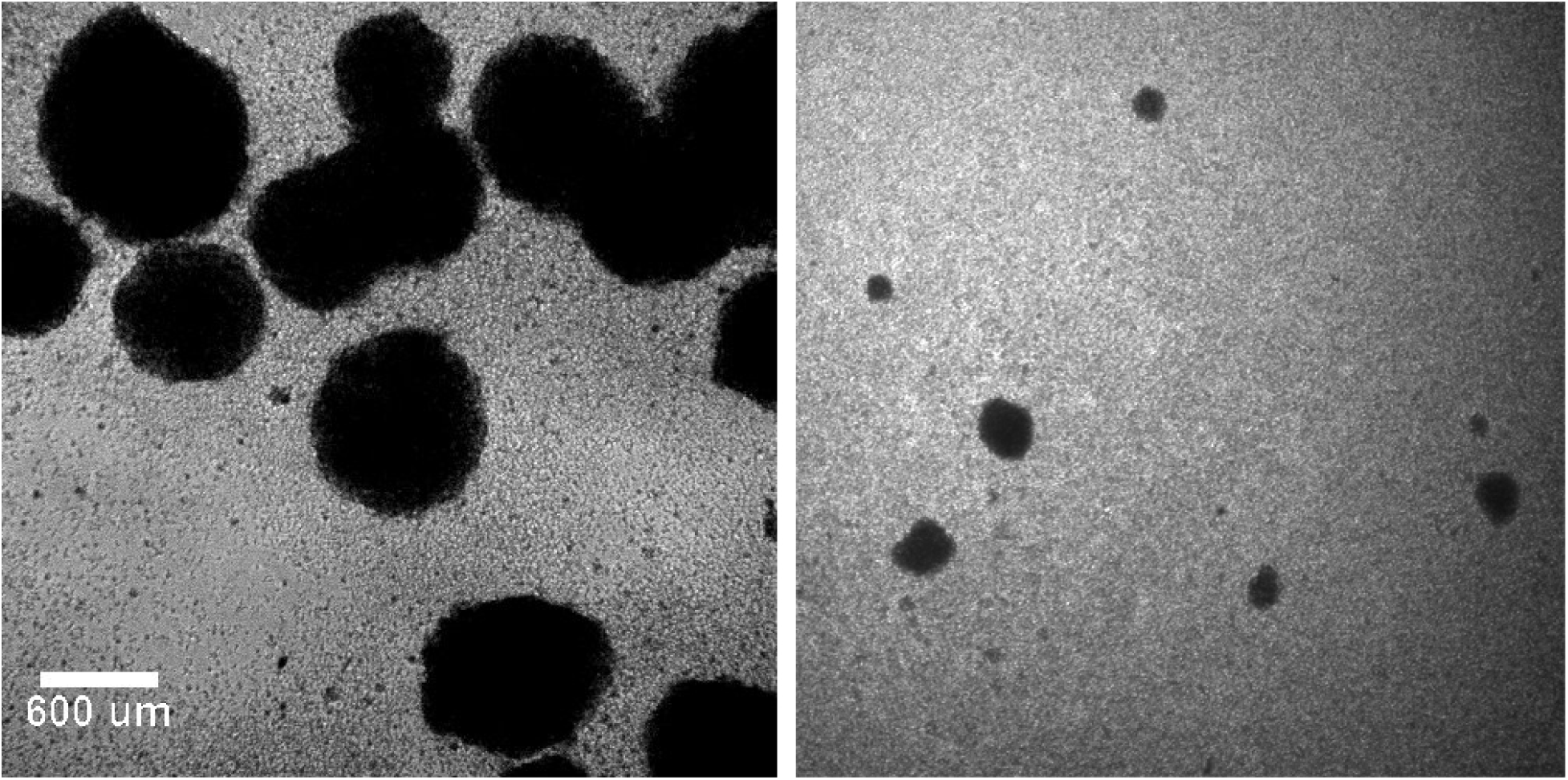
Left: CAR T cells mixed at E:T ratio with RL CD19 KO cells (24 hours after mixing). Right: CAR T cells mixed at same ratio with RL expressing CD19 under LMNB1 promoter (24 hours after mixing).

### Morphological features of CAR T-cell activation and target-cell killing

Live-cell observation showed that CAR T cells displayed amoeboid morphology and high motility, with estimated migration speeds of approximately 5–15 μm/min (Fig. 8, Movie 1).

**Figure 8.**
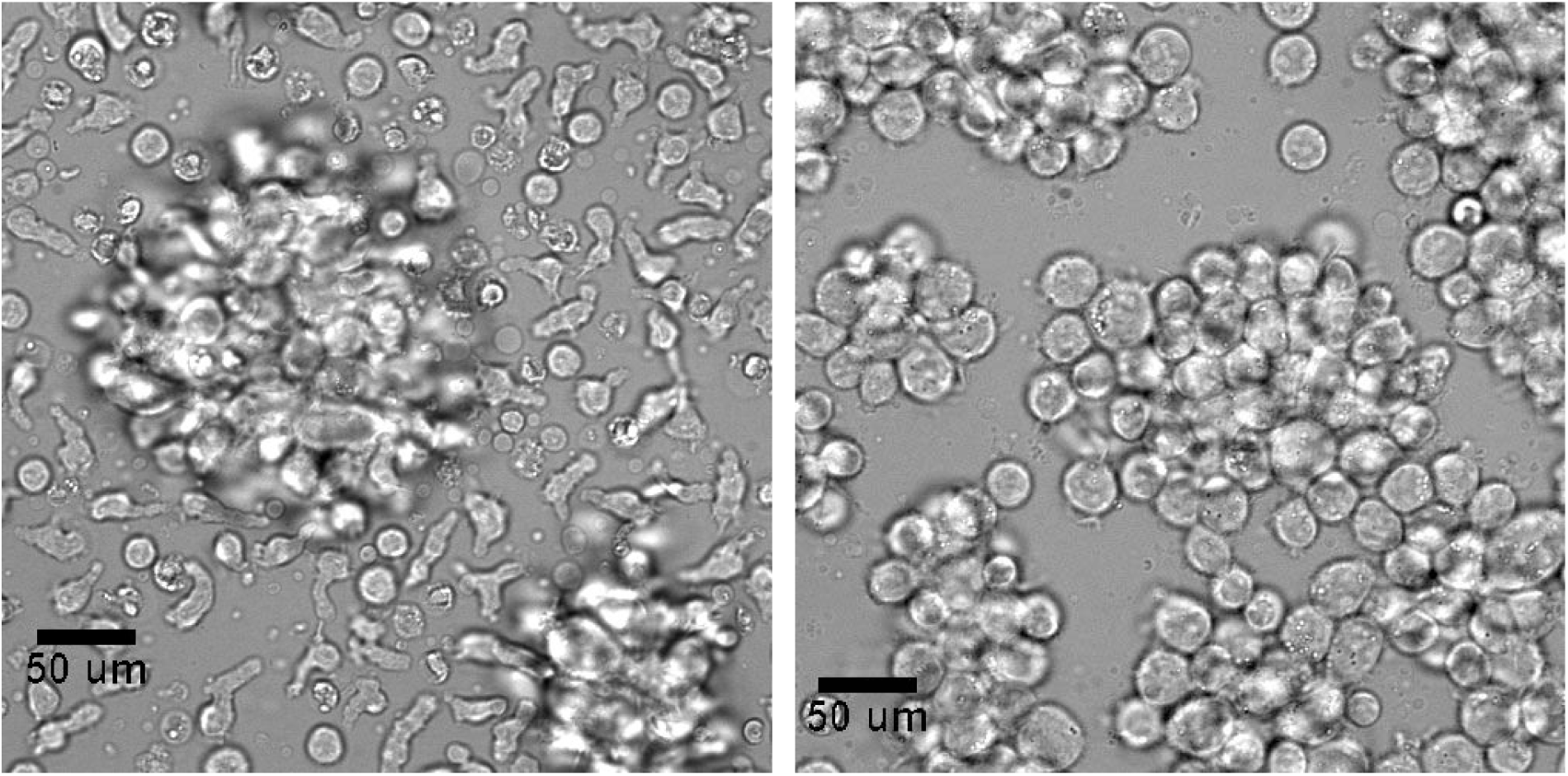
Left: RL cell aggregates and CAR T cells around them. Magnification 40x. Right: RL cell line (ATCC CRL-2261), B lymphoblasts originating from non-Hodgkin’s lymphoma (NHL). These undifferentiated lymphoblasts grow predominantly as large, floating multicellular aggregates in suspension culture. RL cells aggregates look different when compare with RL cells aggregates in wells without CAR T cells (Right). CAR T cells penetrate inside of RL cells aggregates, create mixed RL/CAR T cell aggregates during killing target cells.

T cells are small cells, and they do not look like suspension cells i.e. they are slightly attached to the bottom flask and during migration on flat environment they represent mesenchymal-like phenotype (Tabdanov et al., 2021) i.e. lamellipodium-driven flat adhesive spreading (Saitakis et al., 2017). Small, immobile, spherical CAR T cells were typically observed immediately before cell division, consistent with mitotic rounding that facilitate spindle formation (Fig. 9, Movie 1). In high-density cultures, CAR T cells exhibited both highly motility (cells move with velocities 5-15 μm/min) like phenotypes and low-motility states (0 – 5 μm/min). However, despite observed CAR T adhesion to substrate we observed that even very gentle medium flow can move cells in the flow direction (Movie 3). Thus, CAR T cell lamellipodia that spread on cultureware surface are not firmly attached to the surface and CAR T cells can be easily split by adding new medium, gentle shaking of flack and transferring all flask content to other flask.

**Figure 9.**
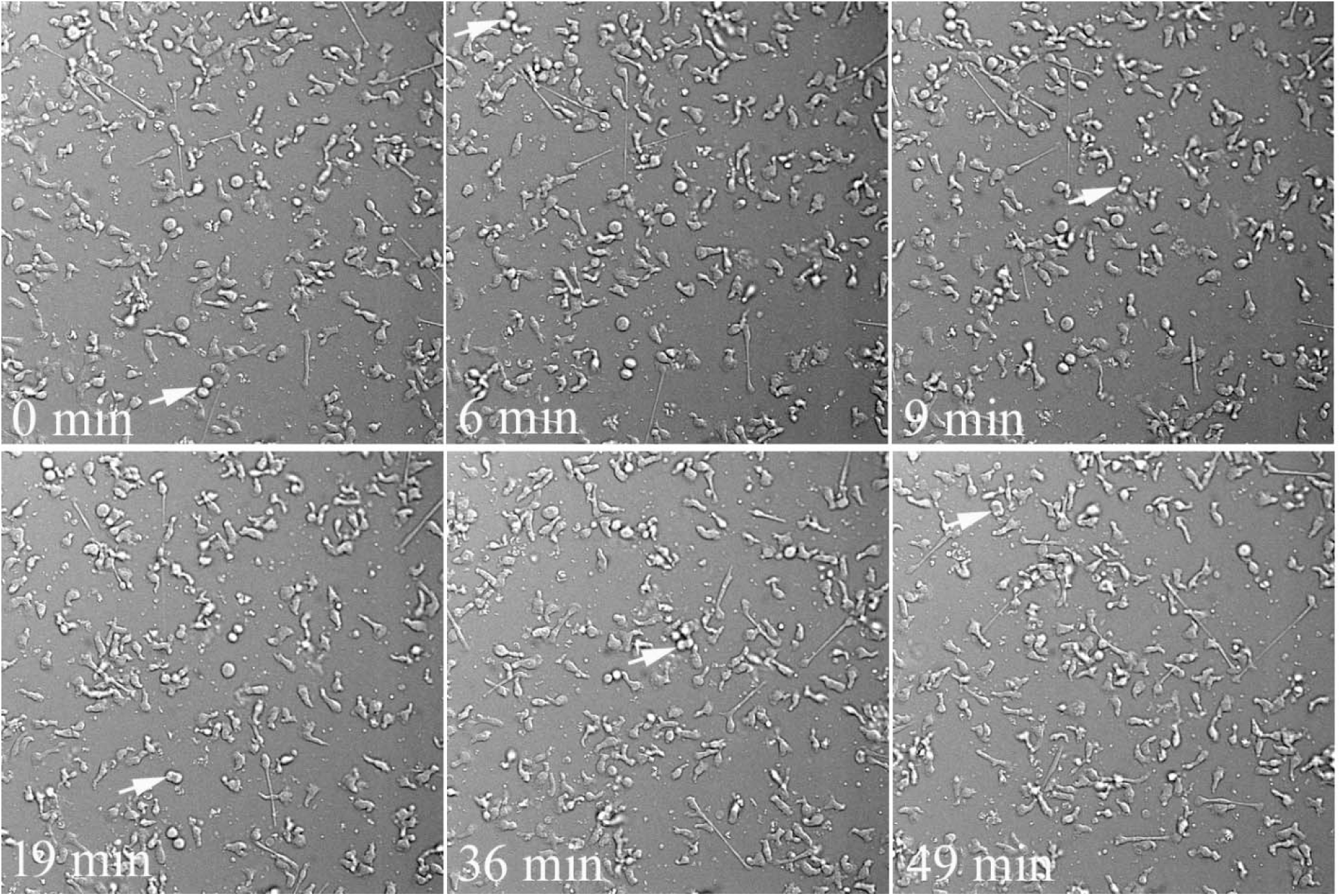
The field of view was taken far away from target cells aggregates. CAR T cells in field of view undergo division events (shown by white arrows). The majority of CAR T cells exhibit high-motility (active movement). Cells that are preparing for division have spherical shape (see Movie 1 for this field of view).

Upon contact with CD19-positive RL targets, CAR T cells formed conjugates and induced morphological changes in target cells consistent with cytolytic death. Target cells initially exhibited membrane blebbing without immediate loss of the fluorescent viability marker BFP and without early DAPI uptake. After bleb rupture, BFP leaked from the target cells (Fig. 10, Movie 2) or, conversely, DAPI entered the cytoplasm and nucleus (Movie 4), indicating loss of membrane integrity. The death process unfolded over several hours and resembled previously described granule-exocytosis-mediated cytotoxic T-cell killing, in which target cells round up, detach, bleb, and subsequently lose membrane integrity (Waterhouse et al., 2006). Similar processes were observed and described in study where cytotoxic T lymphocytes (CTL) induce death of adherent target cells (Waterhouse et al., 2006). To characterize the morphology of death, authors co-incubated CTLs from wild-type mice with MS9II cells at effector:target ratio (2:1). Difference in cells morphology allows easily to identify CTLs (motile and small) from MS9II cells (large, flat, and adherent). Target cell death was detected by time-lapse microscopy and staining with annexin V and propidium iodide (PI). The loss of plasma membrane integrity was detected by the passive uptake of PI (Goldstein et al., 2000). Authors concluded that MS9II cells underwent changes that were virtually identical to those of cells killed by purified perforin and granzyme B. After conjugation, the target cells rapidly rounded and detached from the culture dish. The plasma membrane then became ruffled (blebbed), and, after a further period of time, the cell began to swell.

**Figure 10.**
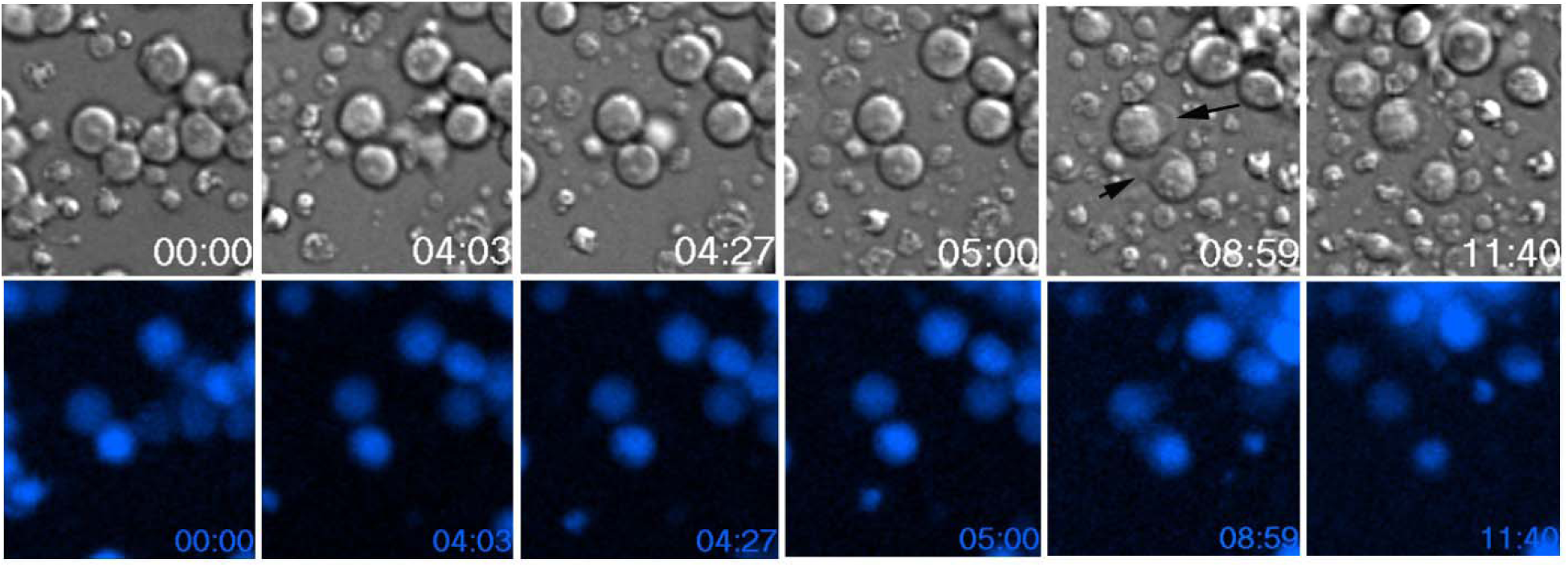
Anti-CD19 CAR-T cells induce cytolisis of CD19 expressing RL cells expressing BFP viability marker (membrane “blebs” shown by black arrows on up DIC panel). Time: hours:minutes. See Movie 2.

### Intermediate CD19 density supports the strongest cytokine release and expansion

Among the RL-derived target-cell panel, the most effective expansion was observed with cells expressing intermediate CD19 density, particularly the RL line in which CD19 was expressed under the LMNB1 promoter. In these cultures, CAR T cells released higher levels of cytokines and expanded more efficiently than when co-cultured with targets expressing either very low or very high CD19 density. We have tested 6 cytokines: INF-γ, IL-10, IL-1β, IL-6, IL-8 and TNF-α. We detected that only four cytokines i.e. INF-γ, TNF-α, IL-10 and IL-8 showed increased release after RL/CAR-T cell mixing (data not shown). Dynamics of INF-γ release when CAR T cells co-incubated with RL expressing different CD19 densities shown in Fig. 11.

**Figure 11.**
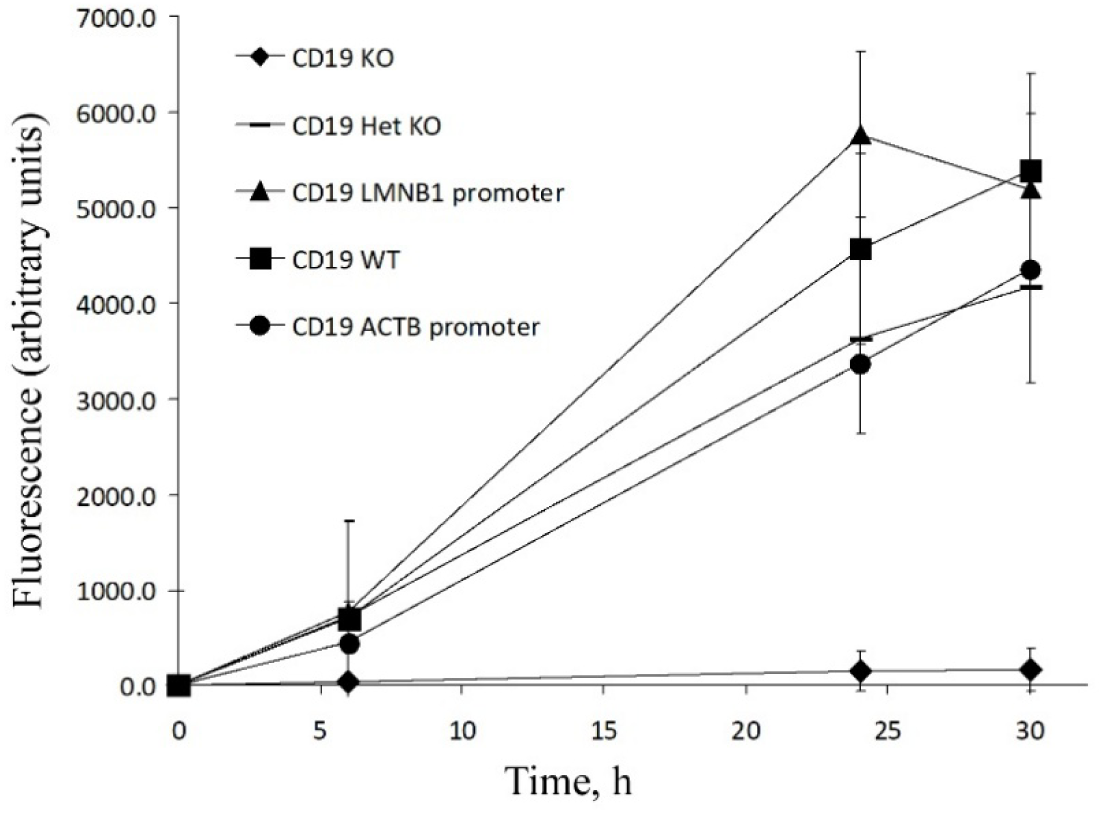
INF-γ release during co-incubation of CAR T cells with RL CD19 lines. A MILLIPLEX Human Cytokine/Chemokine kit (Sigma-Aldrich) was used for testing the control medium and samples.

As CAR T cells lysed the RL target cells, the size of RL aggregates progressively decreased until they disappeared. At that point, the number of CAR T cells increased rapidly, and within 24 hours the cells could form a confluent monolayer on the flask surface (Fig. 12).

**Figure 12.**
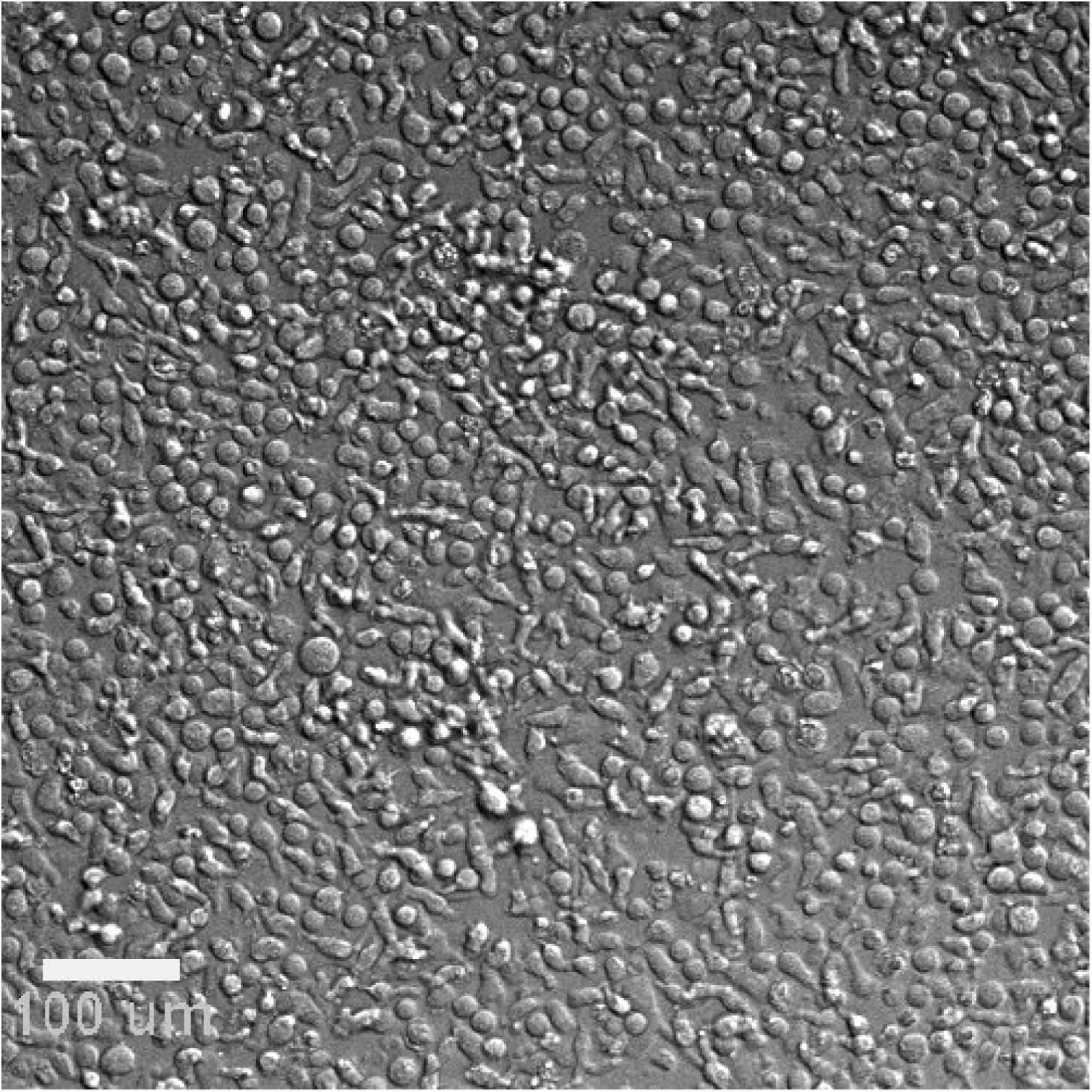
CAR T cells expanded to confluent density after lysis of all RL target cells. 20x magnification.

These findings support the conclusion that intermediate antigen density provides the best balance between productive activation and avoidance of excessive trogocytosis-or exhaustion-associated stress. Cytokine profiling further supported this interpretation. Analysis of control medium and experimental samples showed that the culture medium contained IL-2, whereas four cytokines—IFN-γ, TNF-α, IL-10, and IL-8—were substantially increased after mixing CAR T cells with RL targets. Notably, the highest cytokine release occurred in co-cultures with RL cells expressing intermediate CD19 density under the LMNB1 promoter (Fig. 11), whereas target cells with the highest CD19 density, driven by the ACTB promoter, induced lower cytokine release. As expected, co-culture with CD19-knockout RL cells did not induce substantial cytokine production.

### Sustained expansion depends on repeated antigen exposure

Sustained adCAR-T expansion required continued availability of antigen-positive target cells. Once target cells were eliminated completely, CAR T cells entered a contraction phase consistent with activation-induced cell death (AICD). The first visually-detectable step before AICD is self-aggregation of CAR T cells in absence of target cells (Fig. 13).

**Figure 13.**
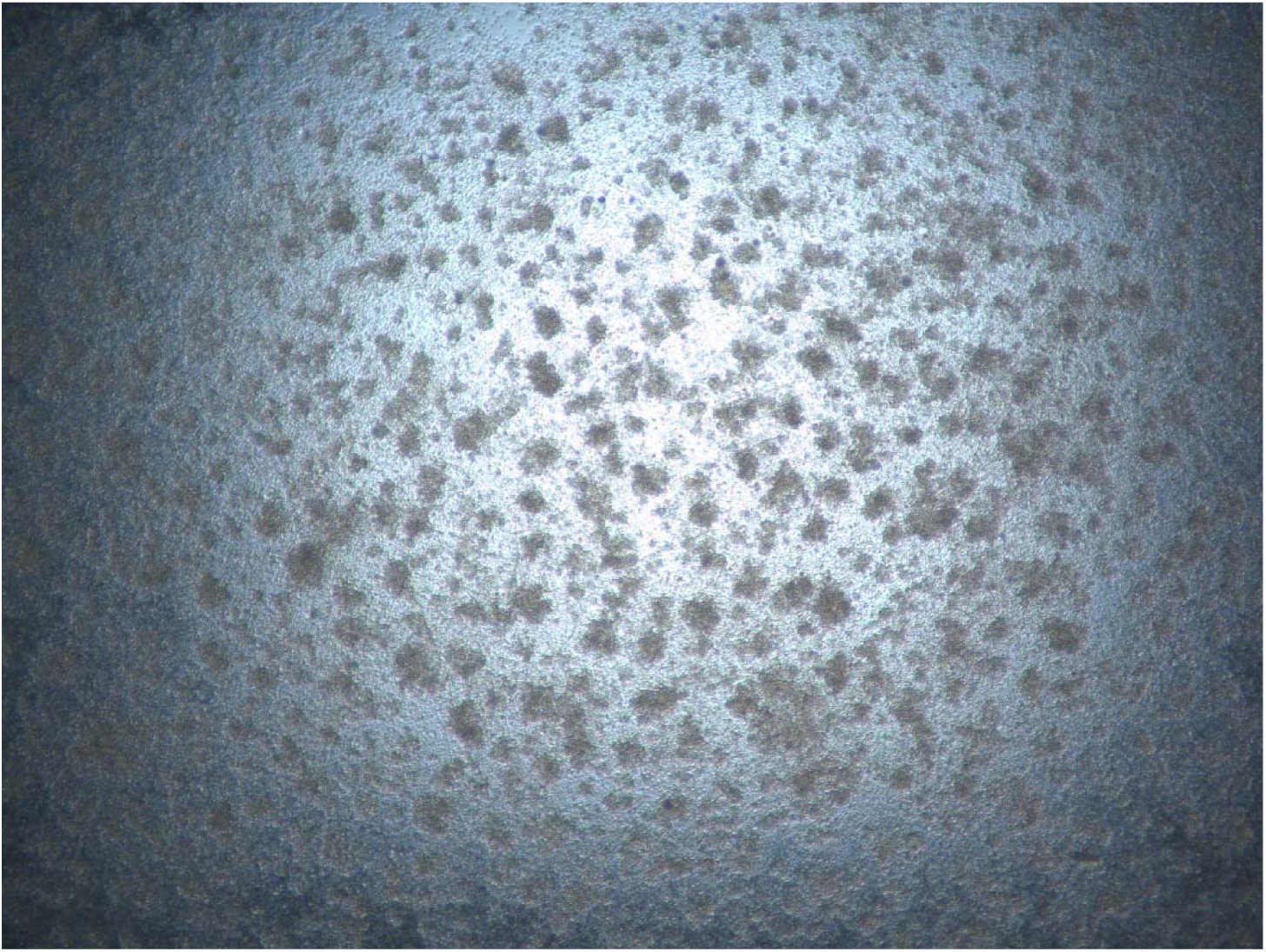
Self-aggregation of CAR T cells during absence of target cells. Magnification 4x.

Activation-induced cell death (AICD) is a mechanism of programmed cell death (apoptosis) in T cells triggered by repeated stimulation of the T-cell receptor (TCR). It serves to remove overactivated or self-reactive T cells, maintaining immune tolerance and homeostasis (Green et al., 2003). Activation-induced cell death (AICD) is a major molecular mechanism that eliminates activated T cells through apoptosis or cell suicide, as part of a process often referred to as ‘clonal elimination (Arakaki et al., 2014). AICD can occur in a cell-autonomous manner and is influenced by the nature of the initial T-cell activation events. AICD primarily occurs via the Fas/FasL death receptor pathway, where Fas ligand (FasL) binds to the Fas receptor (Arakaki et al., 2014). AICD is very important for contraction phase that happened after an infection has been cleared and the number of activated T cells needs to be reduced. In one study authors developed in vitro model system of AICD in which they evaluated the ability of the T helper (Th) cells (CD4+ T cells) to prevent CTL apoptosis due to AICD. The results show that the presence of Th cells significantly reduced CTL susceptibility to AICD through a cell contact-dependent mechanism, indicating a novel method by which Th cells are able to help established CTL responses through increased CTL survival (Kennedy and Celis, 2006). Thus, the effect CD4+ T cells protective function may have on both CTL maintenance and memory. Since CAR T cells prepared for IV injection contain both CD3+ and CD4+ T cells, the last ones may serve as protectors of CD3+ CAR T cells from AICD during eradication of target cells. However, it is possible that this protection function can decline once CD+ CAR T cells stop lysis of target cell. During expansion CD8+ CAR T cells undergo multiple repeated encounters with antigen-bearing target cells. Such multiple encounters and lysis events should cause activation-induced cell death (AICD) of activated CD8+ T cells (Brunner et al., 1995). The mechanism of AICD include Fas(CD95)/FasL interactions required for programmed cell death after T-cell activation (Ju et al., 1995). It was shown that T cells death, but not activation, can be selectively prevented by a soluble Fas-immunoglobulin fusion protein Repeated addition of fresh target cells prevented premature contraction and allowed iterative cycles of killing and re-expansion. In practice, the disappearance of large floating target-cell aggregates served as a convenient visual indicator that a new portion of target cells should be added. The aggregate-forming behavior of RL and Raji-derived lymphoblastoid targets is therefore advantageous for process monitoring, particularly in conjunction with in-incubator live-cell imaging systems.

We found that maintaining an effector-to-target ratio of approximately 2:1 after each addition of target cells may be beneficial, allowing a fraction of CAR T cells to remain engaged in division while others actively lyse targets. As the CAR T cell population expanded to confluence, cultures could be transferred sequentially to larger vessels. For example, a confluent T75 flask could be transferred to a T175 flask, and subsequently to a T500 flask, which may accommodate approximately 80–100 million CAR T cells at confluence.

### Target-cell lysis may contribute metabolic support to the culture environment

During expansion, CAR T-cell-mediated lysis of target cells releases protein-rich cytoplasmic content into the medium. We hypothesize that these products may partially supplement the nutrient environment, although this remains to be tested directly. It was shown that CAR T amoeboid-like motility depends on mitochondrial metabolism, and ATP is known to support both fast-directional migration and slow motility (Ledderose et al., 2018; Simula et al., 2018) of activated T cells and inhibition of its production reduces T cell motility (Haas et al., 2015). It was shown that human CD8^+^ T cell 3D motility is mainly supported by the tricarboxylic acid (TCA) cycle fueled by glucose and glutamine (Simula et al., 2024). This study data indicate that (*i*) an effective CD8^+^ T cell 3D motility is supported mainly by glucose-and glutamine-fueled TCA cycle sustaining both ATP and mtROS production from mitochondria and (*ii*) strategies targeting mitochondrial metabolism are effective in increasing the intratumoral infiltration of CD8^+^ CAR T cells to help them reach and kill tumor cells in preclinical solid tumor models. During acute deprivation (1 h), the absence of glucose, but not glutamine, slightly but significantly reduced 3D motility. After prolonged deprivation (48 h), both nutrients halved 3D motility in a similar way. These data suggest that glucose and glutamine support similarly T cells 3D motility as they can compensate for each other during a short time, while for longer periods both are similarly required, like it was observed for T cell proliferation. Overall, these data indicate that glucose and glutamine, but not fatty acids (FAs,) are required to support CD8^+^ T cell 3D motility and proliferation under normal conditions (Simula et al., 2024). Because activated T-cell motility and proliferation depend strongly on mitochondrial metabolism, ATP generation, and nutrient availability, optimization of glucose and glutamine levels during expansion may improve performance further. In our system, CAR T cells were expanded in a commercial CAR T-cell medium (ProMab Biotechnologies, Inc. Catalog # PM-CAR2000) formulated for high-density growth, but the manufacturer did not specify glucose or glutamine concentrations. Future work should therefore quantify nutrient consumption during adCAR-T expansion and determine whether supplementation further improve proliferation and CAR T “functional fitness”.

### Recommended alternative target-cell panel

Because the RL CD19 panel is not available, we recommend using Raji CD19 antigen panel cell lines that are commercially available (Sigma-Aldrich product # ATG001). These lines span multiple levels of CD19 surface density, from complete knockout to expression substantially above wild-type Raji levels (Fig. 14). Like RL cells, Raji cells grow as large floating multicellular aggregates and reportedly show minimal PD-L1 (B7-H1) induction after IFN-γ stimulation (see Fig. 1A from (Berthon et al., 2010) showing PD-L1 expression in leukemic cell lines and blast cells from Acute Myeloid Leukemia: LAMA-84, HL60, K562, U937, KG1a, THP-1 and lymphoid lines (Raji, Jurkat) with or without incubation with 500 IU/ml IFN-γ for 24 h). These properties making Raji cells suitable for the same expansion logic.

**Figure 14.**
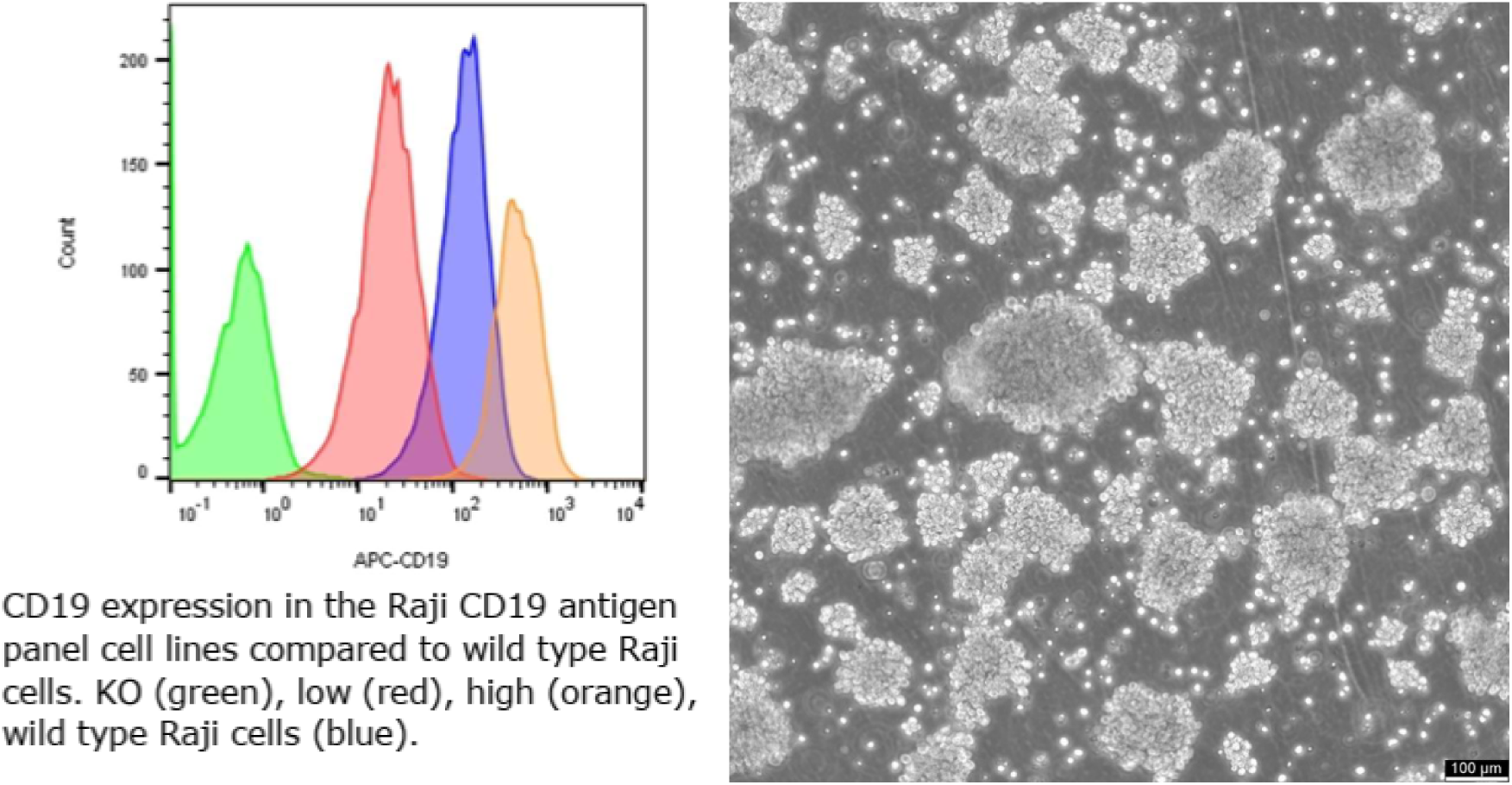
Raji CD19 Antigen Panel Cell Lines (Sigma-Aldrich, ATG001). Raji cell line originated from human Burkitt lymphoma B cells. These undifferentiated lymphoblasts grow predominantly as large, floating multicellular aggregates in suspension culture. Raji cells demonstrate tumorigenicity when xenografted into immunodeficient mice. According to the literature Raji cells do not express PD-L1 even after stimulation with INF-γ (see Fig. 1A from (Berthon et al., 2010).

### Antigen-driven expansion defines a hybrid manufacturing paradigm

Taken together, these findings suggest that CAR T-cell manufacturing can be organized around a third paradigm distinct from both conventional *ex vivo* production and emerging *in vivo* gene-delivery approaches. Standard ex vivo manufacturing offers high control but relies on artificial activation, long culture duration, and carries a risk of exhaustion. *In vivo* CAR T generation simplifies logistics but introduces significant safety concerns related to systemic gene delivery. The adCAR-T approach defines a hybrid alternative: it preserves the control of ex vivo expansion while using physiological, antigen-dependent activation to drive rapid proliferation in a checkpoint-silent setting. In principle, this may reduce exhaustion, shorten manufacturing time, and improve the overall safety profile relative to systemic in vivo gene-delivery strategies. It is known that primary activated cells undergo a programmed burst of about 15–20 cell divisions, with the capacity for at least 3 cell divisions per day (De Boer et al., 2003). This results in massive clonal expansion and is the basis for the specificity and memory that typifies adaptive immunity.

## Discussion

The present study proposes an antigen-driven CAR T-cell expansion platform (adCAR-T) as a hybrid manufacturing paradigm positioned between conventional ex vivo production and emerging in vivo CAR-T generation strategies. The central premise is that freshly transduced CAR T cells should not be expanded under purely artificial conditions, but instead should undergo repeated, controlled encounters with antigen-bearing target cells that mimic the biological context they will face after infusion. In this framework, antigen engagement, target-cell lysis, cytokine release, and iterative proliferation are coupled into a self-reinforcing ex vivo process. We propose that this design may accelerate manufacturing while better preserving the functional properties required for tumor clearance in vivo.

### adCAR-T as an alternative to current manufacturing paradigms

Current CAR-T platforms each solve some problems while creating others. Standard autologous ex vivo manufacturing provides strong process control and well-defined product characterization, but it is slow, labor-intensive, expensive, and biologically artificial. In vivo CAR-T approaches are attractive because they aim to bypass leukapheresis, centralized manufacturing, and prolonged culture, but they shift the major risks toward systemic delivery, biodistribution, off-target transduction, and limited control over how many therapeutic cells are actually generated. The adCAR-T approach is intended as a third option: it preserves ex vivo control over transduction and product handling while replacing artificial stimulation with physiologically relevant antigen-driven activation.

This distinction may be important. In conventional workflows, CAR T cells are activated with anti-CD3/anti-CD28 reagents and expanded for days or weeks before they ever encounter their intended targets. By contrast, in adCAR-T cultures, CAR T cells engage antigen immediately after transduction and expand through repeated cycles of recognition and killing. We hypothesize that this “battlefield” mode of expansion better prepares the cells for the biological demands of tumor elimination after infusion.

### Comparison with Existing Paradigms

Current CAR-T approaches fall into two categories:

1. Ex vivo manufacturing

`o` High control
`o` Artificial activation
`o` Risk of exhaustion
`o` Long production time
2. In vivo generation

`o` Simplified logistics
`o` Reduced manufacturing burden
`o` Safety concerns related to systemic gene delivery via LV vectors and LNPs

The proposed method introduces a third approach:

3. Antigen-driven hybrid expansion

`o` Physiological activation
`o` Controlled environment
`o` Reduced exhaustion
`o` Rapid expansion
`o` Improved safety profile

### Why in vivo CAR-T may face intrinsic quantitative and biological limitations

In vivo CAR-T has generated understandable excitement because it promises a faster, more accessible, and potentially less expensive route to treatment. The review “CAR-T cell manufacturing: Major process parameters and next-generation strategies” discusses current and novel CAR T-cell processing methodologies and the quality control systems needed to meet the increasing clinical demand for these exciting new therapies (Ayala Ceja et al., 2024). The ability of CAR T cells to expand in vivo (T-cell ‘fitness’) is associated with anti-tumor responses and can be impacted by the manufacture method used.

However, several constraints remain. First, the number of CAR T cells generated in vivo after intravenous delivery of lentiviral vectors or mRNA-loaded lipid nanoparticles is likely to be much lower than the number of cells administered in current ex vivo protocols. Even if in vivo delivery yields from several million CAR T cells to tens million, this is still likely to be far below the hundreds of million or up to billions that commonly used in approved therapies.

Second, these newly generated cells are expected to encounter tumor immediately in a setting where the initial effector-to-target ratio is highly unfavorable. In hematologic malignancies such as B-ALL, tumor burden at diagnosis or relapse can be extremely high, with leukemic cells present in bone marrow, blood, and other tissues in very large numbers. Under such conditions, a relatively small in vivo-generated CAR T-cell population may be forced into intense serial killing early, increasing the probability of rapid exhaustion, fratricide following trogocytosis, receptor downregulation, or death before sufficient population-level amplification occurs.

Third, non-integrating in vivo platforms based on mRNA lipid nanoparticles introduce a separate limitation: transient CAR expression. Although transient expression improves genomic safety, it may also mean that T cells that proliferate after initial activation progressively lose CAR surface expression, thereby diluting the functional therapeutic population. Integrating vectors avoid this problem but reintroduce the risks of insertional mutagenesis and off-target transduction. Thus, the perceived simplicity of in vivo CAR-T must be weighed against the fact that it combines uncertain cellular output with delivery-associated safety liabilities.

### Safety advantages of maintaining genetic modification ex vivo

One of the clearest advantages of adCAR-T is that gene transfer remains confined to a controlled ex vivo environment. This allows direct control over transduction conditions, transduction efficiency, target-cell exposure, and product characterization before infusion. By contrast, in vivo approaches depend on systemic delivery of viral or nanoparticle-based constructs and therefore inherently risk unintended gene transfer into non-T-cell populations.

This is not only a theoretical concern. Even during standard ex vivo manufacturing, accidental transduction of a leukemic B cell has been shown to generate CAR-expressing malignant cells capable of escaping recognition through epitope masking (Duncan et al., 2024; Moiani et al., 2012). In a systemic in vivo delivery setting, such events could become even harder to prevent or detect. Likewise, the long-term risk of insertional mutagenesis remains a serious issue for integrating vectors, particularly if administered intravenously. For these reasons, preserving genetic modification within a tightly controlled ex vivo step may represent a major safety advantage.

### The importance of E:T ratio and target-cell design

A major conceptual point emerging from this work is that CAR T-cell expansion is strongly shaped by the starting effector-to-target ratio and by the biological properties of the target cells used for stimulation. In vivo, the starting E:T ratio is effectively uncontrolled and often highly unfavorable. In patients with B-cell acute lymphoblastic leukemia (B-ALL), the number of leukemia cells (lymphoblasts) in the body is generally very high at diagnosis, often numbering in billions or trillions. The bone marrow is typically packed with >20 % and sometimes nearly 100% leukemia cells, which then circulate in the blood and spread to other organs (Chiaretti et al., 2014; Terwilliger and Abdul-Hay, 2017). Thus, after IV injection of 250 × 10^6^ CAR T cells in patient the start E:T ratio will be in range 1:400 to 1:4000. The injection of one billion CAR T cells can move E:T ratio to more favorable range. However, with trillion cancer cells in patient blood the injection of one billion CAR T cells will allow to reach E:T ratio 1:1000.

In adCAR-T cultures, by contrast, antigen density, E:T ratio, and checkpoint-ligand expression can all be deliberately selected and adjusted over time. We propose that this controllability is central to the performance of the system. If the E:T ratio is too low, each CAR T cell may be forced into excessive serial killing, resulting in rapid granule depletion, exhaustion, trogocytosis-associated fratricide, or activation-induced cell death. If target cells express excessive levels of antigen, overly stable synapses may again increase trogocytosis and functional collapse. Conversely, if antigen density is too low, activation becomes insufficient. Our data suggests that intermediate antigen density provides the best balance, supporting strong cytokine release and rapid proliferation without driving excessive dysfunction. The checkpoint profile of the target cells is also very important. The development of CAR-T cell exhaustion has been proposed to be triggered by co-inhibitory pathways such as PD1/PD-L1 (Yin et al., 2018). IFN-γ released by activated CAR T cells can induce PD-L1 on many tumor cells, turning a useful activation signal into a self-limiting suppressive loop. By selecting target cells with low basal PD-L1 that do not substantially upregulate PD-L1 after IFN-γ exposure, the adCAR-T platform avoids this negative feedback during expansion. We consider this a key design principle rather than a minor technical optimization.

### Potential implications for lymphodepletion

Lymphodepleting chemotherapy remains standard in current CAR-T therapy because it creates space for infused cells, raises homeostatic cytokine levels, and reduces competition and suppression from endogenous lymphocyte populations. Whether adCAR-T products could reduce the need for lymphodepletion remains unknown, but the question is worth considering.

Because adCAR-T cells are already activated and expanding before infusion, they may enter the patient in a more competitive and functionally primed state than cells produced by standard protocols. This could, in principle, lessen reliance on host preconditioning. On the other hand, lymphodepletion also reduces regulatory and competing immune populations and may still be essential, especially in patients with high tumor burden. At present, the most realistic position is that lymphodepletion will probably still be needed in many settings, but perhaps not always at the same intensity. This will require direct experimental and clinical testing.

### Functional state, persistence, and exhaustion

A central unresolved issue in CAR-T therapy is how to generate cells that are not merely numerous, but also durable and capable of persistence after infusion. Clinical relapses often reflect either inadequate persistence of otherwise functional CAR T cells or tumor escape through antigen modulation therapeutic interventions for CRS on the durability of CAR remission remains unknown (see Figure 1 in (Shah and Fry, 2019). We hypothesize that adCAR-T may be particularly relevant to the first category.

In the proposed system, CAR T cells expand while repeatedly engaging antigen and killing target cells. This may select for cells that are both highly functional and able to sustain repeated cytotoxic encounters. In principle, such cells may be better prepared for continued activity in the patient than cells expanded under bead-driven conditions. We did not observe obvious signs of overt exhaustion during the expansion phase, although this will need formal validation using phenotypic, transcriptional, and functional assays.

An important future question is the balance of CD8+ and CD4+ CAR T cells before and after adCAR-T expansion. CD8+ cells likely execute most direct cytolysis, whereas CD4+ cells may provide critical proliferative and survival support through cytokine secretion and possibly protection against activation-induced apoptosis (Bove et al., 2023). It will be informative to determine whether antigen-driven expansion shifts this balance and whether that shift contributes to persistence or high anti-tumor activity.

Related to this, asymmetric cell division (ACD) and c-Myc partitioning may be highly relevant to the final product state. ACD − the partitioning of cellular components in response to polarizing cues during mitosis—plays roles in differentiation and development. T lymphocytes, upon activation by antigen-presenting cells (APC), can undergo ACD, wherein the daughter cell proximal to the APC is more likely to differentiate into an effector-like T cell and the distal daughter more likely to differentiate into a memory-like T cell (Chang et al., 2007). In the case of activated T cells, one daughter cell became the rapidly dividing effector T cells that launch the immediate attack on infectious agents and other threats (Verbist et al., 2016). The other daughter cell became the slowly dividing memory T cells that function like sentries to provide long-term protection against recurring threats. One study showed that the regulatory protein c-Myc distributes during asymmetric cell division directly influences the fate and roles of activated T cells - during asymmetric cell division of activated T cells, high levels of c-Myc accumulated in one daughter cell. The daughter cells with low levels of c-Myc functioned like memory T cells, proliferating to mount an immune response a month later when mice were again exposed to the virus. It was shown that T cell proliferation depends on c-Myc expression level in daughter cells. The c-Myc^high^ T cells divided nearly ten times the frequencies of c-Myc^low^ T cells. In addition, sorted c-Myc^high^ T cells continue to proliferate much more rapidly than c-Myc^low^ T cells when placed back in culture for 48 hours.

If adCAR-T cultures preserve or generate a mixture of rapidly dividing effector-like cells and more slowly dividing memory-like cells, this could provide a mechanistic basis for both immediate cytotoxicity and long-term persistence. This remains speculative, but it is a promising direction for future analysis. It is known that CAR T cell persistence correlates with improved outcome in cancer patients (Wittibschlager et al., 2023). For example, lymphoma relapses were less frequent among patients with CAR T-cell persistence (29% vs. 60%, *p* = 0.0336), and CAR T-cell persistence in the PB at 6 months was associated with longer progression-free survival (PFS) (Wittibschlager et al., 2023). The observations of CAR T cells propagation at the presence of target cells allow us to conclude that the population maintain persistence during all period without any signs of exhausting.

### Manufacturing implications and practical feasibility

The main practical value of adCAR-T may not simply be that it shortens culture time, although that is important. Rather, its more significant advantage may be that it enables rapid production of a highly active CAR T-cell population from relatively small starting numbers of patient T cells (1-2 million or even less). This could be particularly valuable for heavily pretreated patients who yield poor leukapheresis products and are often excluded from current manufacturing paradigms.

If adCAR-T can reliably start from very small inputs and still generate clinically relevant doses within 2–3 days or even one week, it could lower the threshold for patient eligibility, reduce vector usage, eliminate the need for bead-based activation, and lessen dependence on long and costly bioreactor-based expansion phases. Because lentiviral vector is one of major cost drivers in CAR-T manufacturing (Abdo et al., 2025; Mallapaty, 2024), reducing the number of cells that must be transduced initially could substantially lower per-patient cost.

The culture format itself may also matter. Aggregate-forming target cells are particularly useful because their disappearance provides a simple visual indicator that the next round of stimulation should begin. T-flasks proved practical for early-stage work because target-cell aggregates could be monitored even at low magnification. CAR T cells in these cultures did not behave like classical suspension cells; rather, they displayed weak substrate attachment and active motility, making them easy to transfer during expansion without harsh detachment procedures like trypsinization.

### Constraints and unanswered questions

This study also has clear limitations. The proposed method still requires formal optimization of target-cell inactivation, control for residual target cells (they must be mitotically inactivated via γ-irradiation and labelled by fluorescent viability marker), nutrient management, and product purification before infusion. The possibility of target-cell contamination must be carefully addressed in any translational version of the process. Mitotically inactivated via γ-irradiation K562 cells used for expansion of Natural Killer (NK) cells (Lapteva et al., 2012). Irradiation (commonly 50–100 Gy) permanently arrests K562 growth, ensuring they do not contaminate the final expanded NK cell product that used in human clinical trials. Flow cytometry-based assays should be validated and used to show that γ-irradiated target cells do not proliferate. Constantly expressed viability marker (BFP or other fluorescent proteins) is useful for confirming the elimination of target cells. However, there may be concern that BFP released from target cells can induce an immune response in the patient after infusion. CAR T purification process includes multiple, critical wash cycles to ensure product purity, safety, and efficacy (Levine et al., 2017). These cycles eliminate cell debris, cytokines, medium serum and any other proteins that were released during manufacturing process. Thus, viability protein most likely will be completely eliminated.

In addition, although our observations support rapid expansion and sustained activity, the present work does not yet define the transcriptional, metabolic, or epigenetic state of the final product.

Several specific questions should be prioritized in future work:

1. What are the optimal antigen density and E:T ratio ranges for different CAR designs?
2. How does adCAR-T expansion affect CD4/CD8 composition, memory phenotype, and exhaustion markers?
3. Does the method generate cells with improved persistence after infusion relative to bead-expanded controls?
4. Can residual target-cell contamination be eliminated robustly and reproducibly?
5. Will the method work equally well for allogeneic CAR-T products?

The final question is particularly interesting. In principle, there is no obvious reason why allogeneic CAR T cells could not be expanded using the same antigen-driven logic. If so, adCAR-T might be compatible not only with autologous manufacturing but also with off-the-shelf platforms.

## Conclusion

In summary, we propose antigen-driven CAR T-cell expansion as a hybrid manufacturing strategy that combines the safety and control of ex vivo production with the biological logic of in vivo CAR T method. In this model, freshly transduced CAR T cells are expanded through repeated, controlled encounters with engineered target cells under conditions in which antigen density, checkpoint signaling, and E:T ratio can be optimized. We hypothesize that this approach may accelerate manufacturing, reduce cost, lower input-cell requirements, and generate CAR T cells that better suited for persistence and function after infusion. Proposed adCAR-T method changed time for cell production from 5-14 days (for most FDA approved CART-cell products, see Figure 1 from (Roddie et al., 2019) and Figure 1 from (Martinez-Cibrian et al., 2022) to 2-3 days. But it is not main advantage of this method. Main advantage is that CAR T cells obtained by this method are 100% active for vigorous attacking and lysis of cancer cells that express CAR-targeted antigen. After infusion they will continue to attack and lyse patient cancer cells. CAR T cells exhausting in patient body is not excluded especially if cancer cells will start to express PD-L1 after CAR T release high concentration of INF-γ. As an option, patient cancer cells could be tested in advance for possible INF-γ induced PD-L1 expression and if it is the case CAR T infusion could be combined with infusion of anti-PD-1 antibodies (Ouyang et al., 2024).

Lymphodepleting chemotherapy is usually required for blood cancer patients. Most likely that it will be required for CAR T obtained by adCAR-T method. Cytokine release syndrome (CRS) is also possible. Interleukin-6 (IL-6) is recognized as the most critical cytokine in the pathophysiology and management of Cytokine Release Syndrome (CRS), particularly in CAR T-cell therapy. Elevated serum IL-6 levels are strongly correlated with CRS severity, driving vascular leakage, fever, and acute-phase reactions (Lee et al., 2014). Therefore, CRS treatment focuses on managing symptoms and controlling the immune response, often with the IL-6 receptor antagonist tocilizumab and corticosteroids. Surprisingly, we did not detect any significant IL-6 release by CAR T cells expanding via adCAR-T method (data is not shown). However, it is known that IFN-γ released early by CAR T cells can initiate the cascade by activating macrophages to produce IL-6 and TNF-α (Imran et al., 2025).

The broader implication is that the future of CAR-T therapy may not be determined by a single winning platform, but by parallel evolution of standard ex vivo, allogeneic, and in vivo strategies. The adCAR-T concept offers a distinct path within this landscape: not fully in vivo, not conventionally ex vivo, but a controlled intermediate state designed to capture the advantages of both. Whether this approach can improve persistence, reduce CD19-positive relapse, and expand access for patients with limited starting material will require direct testing, but the concept provides a plausible new framework for rethinking CAR-T manufacturing.

Supplement movies 1-4 is available at https://www.samdolite.com/movies

## Notes

https://www.samdolite.com/movies

### Competing Interest Statement

The authors have declared no competing interest.

